# The facultative intracellular symbiont *Lariskella* is neutral for lifetime fitness and spreads through cytoplasmic incompatibility in the leaffooted bug, *Leptoglossus zonatus*

**DOI:** 10.1101/2025.01.29.635435

**Authors:** Edwin F. Umanzor, Suzanne E. Kelly, Alison Ravenscraft, Yu Matsuura, Martha S. Hunter

**Affiliations:** Entomology & Insect Science Graduate Interdisciplinary Program, The University of Arizona, Tucson, AZ, United States; Department of Entomology, The University of Arizona, Tucson, AZ, United States; Department of Biology, The University of Texas at Arlington, Arlington, TX, United States; Tropical Biosphere Research Center, The University of the Ryukyus, Japan

## Abstract

The maternally-inherited, intracellular bacterium *Lariskella* (Alphaproteobacteria: Midichloreaceae) has been widely detected in arthropods including true bugs, beetles, a wasp, a moth, and pathogen-vectoring fleas and ticks. Despite its prevalence, its role in the biology of its hosts has been unknown. We set out to determine the role of this symbiont in the leaffooted bug, *Leptoglossus zonatus* (Hempitera: Coreidae). To examine the effects of *Lariskella* on bug performance and reproduction as well as in possible interactions with the bug’s obligate nutritional symbiont, *Caballeronia*, bugs were reared in a factorial experiment with both *Lariskella* and *Caballeronia* positive and negative treatments. Lifetime survival analysis (∼120 days) showed significant developmental delays and decrease in survival for bugs that lacked *Caballeronia,* and *Caballeronia*-free bugs did not reproduce. However, among the *Caballeronia* carrying treatments, there were no significant differences in lifetime survival or reproduction in treatments with and without *Lariskella,* suggesting this symbiont is neutral for overall bug fitness. To test for reproductive manipulation, crossing among *Lariskella-*positive and negative individuals was performed. When *Lariskella-*negative females were mated with *Lariskella* positive males, fewer eggs survived early embryogenesis, consistent with a cytoplasmic incompatibility (CI) phenotype. Wild *L. zonatus* from California and Arizona showed high but not fixed *Lariskella* infection rates. Within individuals, *Lariskella* titer was low during early development (1^st^-3^rd^ instar), followed by an increase that coincided with development of reproductive tissues. Our results reveal *Lariskella* to be among a growing number of microbial symbionts that cause CI, a phenotype that increases the relative fitness of females harboring the symbiont. Understanding the mechanism of how *Lariskella* manipulates reproduction can provide insights into the evolution of reproductive manipulators and may eventually provide tools for management of hosts of *Lariskella*, including pathogen-vectoring ticks and fleas.

**Author Summary:** Arthropods are a highly diverse group of animals with ancient relationships with bacteria. Some maternally-inherited, intracellular bacterial symbionts have evolved strategies to manipulate arthropod host reproduction, ensuring their own spread and persistence. One such strategy is cytoplasmic incompatibility (CI), where symbionts in males prevent viable offspring when these males mate with uninfected females. However, reproduction is ‘rescued’ in crosses with both mates infected, promoting bacterial transmission within host populations. In the leaffooted bug *Leptoglossus zonatus*, crossing experiments revealed a significant reduction in egg survival rates when *Lariskell*a-negative females mated with *Lariskella*-positive males, a typical CI phenotype discovered anew for this symbiont. Additional experiments included measuring performance and fitness of bugs with combinations of *Lariskella* and *Caballeronia* (the primary nutritional symbiont of *L. zonatus*). While *Caballeronia*-negative treatments caused developmental delays and mortality, the presence of *Lariskella* had no effect on development or reproduction. *Lariskella* has been frequently found in pathogen-vectoring arthropods, but its specific role remained unclear. Understanding the role that *Lariskella* plays in *L. zonatus* can potentially help us find future pest control measures against blood-sucking pathogen-vectors by exploiting novel CI mechanism to suppress host populations and hence the transmission of human diseases.

## Introduction

Arthropods harbor a multitude of microbial symbionts with diverse roles. Among those that are most consequential for the biology of their host, many are maternally (vertically) inherited and have relationships that have endured for remarkably long periods of time (1–3). Theory predicts that strictly vertically transmitted symbionts can proliferate through generations if they cause hosts to produce more daughters than do uninfected hosts (4). Some bacterial symbionts achieve this by increasing the total number of offspring, supplying essential nutrients that are deficient in the host’s diet or protecting against parasitism or environmental stress (5–9). However, in some instances, symbionts manipulate host reproduction towards female fitness in ways that solely enhance their own transmission (10–12). Reproductive manipulator symbionts can manipulate their host reproduction in many ways, of which the most common is called cytoplasmic incompatibility (CI) (13–15).

CI symbionts modify host males such that reproduction is sabotaged when infected males with modification factors mate with females lacking the CI symbiont, resulting in few or no offspring. The modification is “rescued” when a modified male mates with an infected female carrying rescue factors (15–18). In the well-studied CI-causing symbiont *Wolbachia* (10,19–21), two cytoplasmic incompatibility genes, *cifA* and *cifB*, have been identified (15,17,18). While there is ongoing debate about the appropriate nomenclature and CI mechanism (22,23), transgenic studies in *Drosophila* suggest a “two-by-one” model where both *cifA* and *cifB* from the *Wolbachia* strain *w*Mel must be expressed in males to induce CI, whereas only *cifA* needs to be expressed in females to rescue CI (24). An alternative hypothesis, known as the toxin-antidote (TA) model, suggests that while both *cif* factors colocalize in germ cells, *cifB* travels with mature spermatids and acts as a toxin to developing embryos unless the corresponding antidote *cifA* factor is present, binding to and neutralizing the toxin’s effects (25,26).

The exact molecular mechanism behind CI is unknown and may vary depending on the host and symbiont strain (24,26). However, the consequence of CI is a clear fitness decrease for symbiont-free females, which are only able to mate successfully with other symbiont-free males, and a relative fitness benefit for females that harbor the CI symbiont and can successfully mate with both symbiotic and aposymbiotic males. Reproductive manipulation can wield significant influence on host ecology and evolution due to rapid changes in host population structure and potentially contribute to speciation by altering mating outcomes and reproductive isolation mechanisms (27–30). Leveraging these manipulation mechanisms can also lead to development of novel pest control strategies (31–34) as well as techniques to limit the spread of deadly disease vectors (35–38).

The focal symbiont of this study, *Lariskella,* is a maternally inherited alphaproteobacterium belonging to the recently characterized family of uncultivable intracellular symbionts in the order Rickettsiales, Midichloriaceae (39,40). Midichloriaceae is an ancestrally aquatic clade of endosymbionts that includes the tick-associated clade *Midichloria* and the arthropod associated clade *Lariskella*, along with other lineages found in amoebae, corals, sponges, and aquatic invertebrates (40). However, Midichloriaceae has received considerably less attention relative to the other lineages of Rickettsiales.

*Lariskella*, provisionally named “*Montezuma*” was first detected in the southern Khabarovsk Territory in Russia from blood and tissue samples from humans experiencing acute fever following tick bites (41). Phylogenetic analysis placed the bacterial 16S rRNA on a distinct branch within the Rickettsiales, with 97% of *Ixodes persulcatus* and 5% of *Haemophysalis concinnae* ticks testing positive for *Lariskella* (41). In *I. persulcatus, Lariskella* is highly prevalent in females (up to 90%) but less so in males (around 30%). This sex-specific distribution is consistent across Russian and Japanese tick populations, suggesting that *Lariskella* may influence reproductive processes or fitness in ticks (42,43). *Lariskella* was found at varying abundances within flea-associated bacterial communities (44), and in the hen flea *Ceratophyllus gallinae*, *Lariskella* is among the dominant bacterial associates (45).

*“Montezuma”* was later found and characterized in seed bugs of the genus *Nysius* (Hemiptera: Lygaeidae) and formally proposed as Ca. *Lariskella arthropodarum* (46). In a survey of 191 species of bugs in the infraorder Pentatomomorpha, *Lariskella* was found in 16 host species, with the highest infection frequencies found in *Nysius* spp (Lygaeidae), at 77%-100% (46). Since then, *Lariskella* has been identified in several other insect orders. In Hemiptera, *Lariskella* was found sporadically in microbiome sequencing data from the lygaeid bug, *Henestaris halophilus* (47) and was found in *Macrosteles maculosus* leafhoppers, where it may contribute to nutrition or host fitness (48). In Coleoptera, *Lariskella* was detected in the myrmecophile beetle of the genus *Cephaloplectus* (Ptiliidae) (49). In weevils in the genus *Curculio*, *Lariskella* exhibits a complex evolutionary history, as their sequences do not align with host phylogenies nor form a monophyletic group, indicating likely horizontal transmission events (50). Additionally, *Lariskella* has also been identified in the tortricid moth, *Epinotia ramella* (NCBI: 3066224) and the chrysidid wasp, *Hedychridium roseum* (NCBI: 3077949) through metagenome sequencing (51). The function of *Lariskella* in all these hosts is unknown. Its potential role as a nutritional endosymbiont in ticks, aiding in the synthesis of essential nutrients deficient in their blood-based diet has been considered (52), but incomplete vitamin biosynthesis pathways observed in genomic analyses suggest a more complex role that requires further investigation (52).

Here we investigated the role of *Lariskella* in the seed-feeding leaffooted bug *Leptoglossus zonatus* (Hemiptera: Coreidae). First, we determined the effects of *Lariskella* on the lifetime fitness of its host in a factorial design, comparing bugs with and without both *Lariskella* and the primary symbiont in this system, the obligate, environmentally-acquired symbiotic gut bacterium *Caballeronia* (Betaproteobacteria: Burkholderiaceae) (53). We tested the hypothesis that *Lariskella* might provide nutritional benefits and rescue host development when *Caballeronia* was absent. In a second experiment, crosses between *Lariskella*-positive and negative adults were performed to investigate whether *Lariskella* caused CI. We found virtually no fitness costs or benefits of *Lariskella,* nor any interaction of *Lariskella* with *Caballeronia.* We also found a pattern of offspring production that suggests *Lariskella* causes CI, providing evidence to add *Lariskella* to the growing list of bacterial symbionts that cause this reproductive manipulation. Lastly, we found high frequencies of *Lariskella* in field populations of *L. zonatus,* as would be expected for a bacterium that causes moderate CI, has a near-perfect rate of maternal transmission, and imposes no fitness costs over the lifetime of the bug.

### Study system

*Leptoglossus zonatus* is a polyphagous agricultural pest widely distributed in the southern and southwestern United States and in South America (54,55). In the Southwest, *L. zonatus* is arboreal and commonly feeds on pomegranate, almonds, pistachio and oranges (56,57). As in *Riptortus pedestris* (Alydidae), the model system for bug-*Caballeronia* interactions, second instar *Leptoglossus zonatus* nymphs acquire their obligate symbiont, *Caballeronia,* orally from a complex assemblage of soil microbes (58,59). In these insects, a very narrow tube (the “constricted region” (CR)) joins the midgut 3rd (M3) and 4^th^ (M4) sections. The CR acts as a sorting organ that allows only specific lineages of bacteria to pass and colonize the M4, which then functions as a symbiotic organ (60,61). In *L. zonatus, Caballeronia* acquisition appears to be obligate for normal development (53). Nymphs that failed to acquire *Caballeronia* experienced developmental delay, high rates of juvenile mortality, and were half the weight of their symbiotic counterparts (53). In *R. pedestris,* genomic and transcriptomic analyses revealed that *Caballeronia* can provide essential amino acids and B vitamins and help recycle metabolic waste, while receiving diverse sugars and sulfur compounds from the host (62).

## Results

### *Lariskella* maternal transmission rate and abundance through development

*Lariskella* was maternally transmitted with >99% efficiency. All the eggs tested (113) were positive for *Lariskella.* The titer of *Lariskella* remained similar throughout the first three developmental stages (1^st^-3^rd^ instar), with an average of 1.95 x 10^4^ copies per nymph. *Lariskella* titer increased after the third instar (Fig. 1a) and was also high in reproductive tissue (testes and ovaries), with an average total abundance similar to the whole-body 4th instar nymph (Fig. 1b). On average, *Lariskella* was similarly abundant in ovaries and testes, although ovaries had higher variance (Fig 1b).

**Fig. 1.**
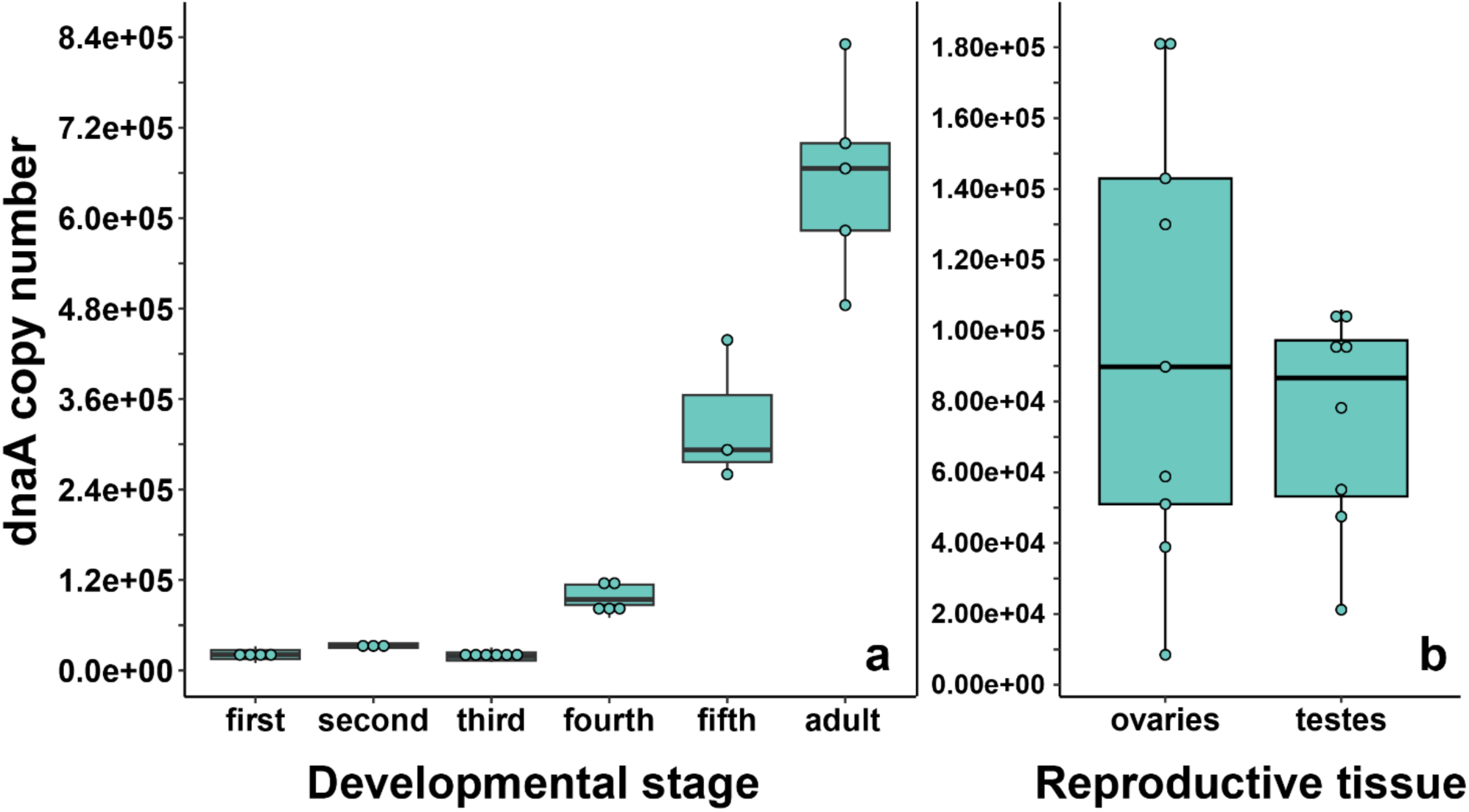
**a)** Absolute *Lariskella* dnaA copy number per individual throughout development (1^st^ instar nymph - adult) and **b)** in reproductive tissue of adults. Each point in 2b represents the paired ovaries or testes from one individual.

### Reproductive tissue and whole-body 16S Illumina sequencing

Amplicon sequencing with universal 16S rRNA primers of whole-body 4^th^ instar nymphs and reproductive tissues was used to characterize the bacteria associated with *L. zonatus.* Unsurprisingly, the dominant sequence variant (SV) in whole-body samples was the obligate nutritional gut symbiont *Caballeronia* (Fig. 2). *Lariskella* reads occurred in low abundance in 3/5 of 4^th^ instar nymphs, with the common gut bacterium, *Enterococcus,* also being abundant in samples without *Lariskella*. In contrast, *Lariskella* was abundant in both ovaries and testes, consistent with a CI-causing phenotype. *Lariskella* was the most consistently present and abundant intracellular taxon found. Importantly, symbionts known to cause CI (*Wolbachia, Cardinium, Rickettsiella, Spiroplasma, Mesenet* and *Rickettsia*) were all absent. *Enterococcus* was also abundant in several reproductive tissue samples. This bacterium is a common gut inhabitant (63,64) and was likely a contaminant from gut disruption during dissections. The genus *Serratia* includes opportunistic pathogens and intracellular symbionts, but this lineage was found in a minority of reproductive tissue samples (43%) (65–67).

**Fig. 2.**
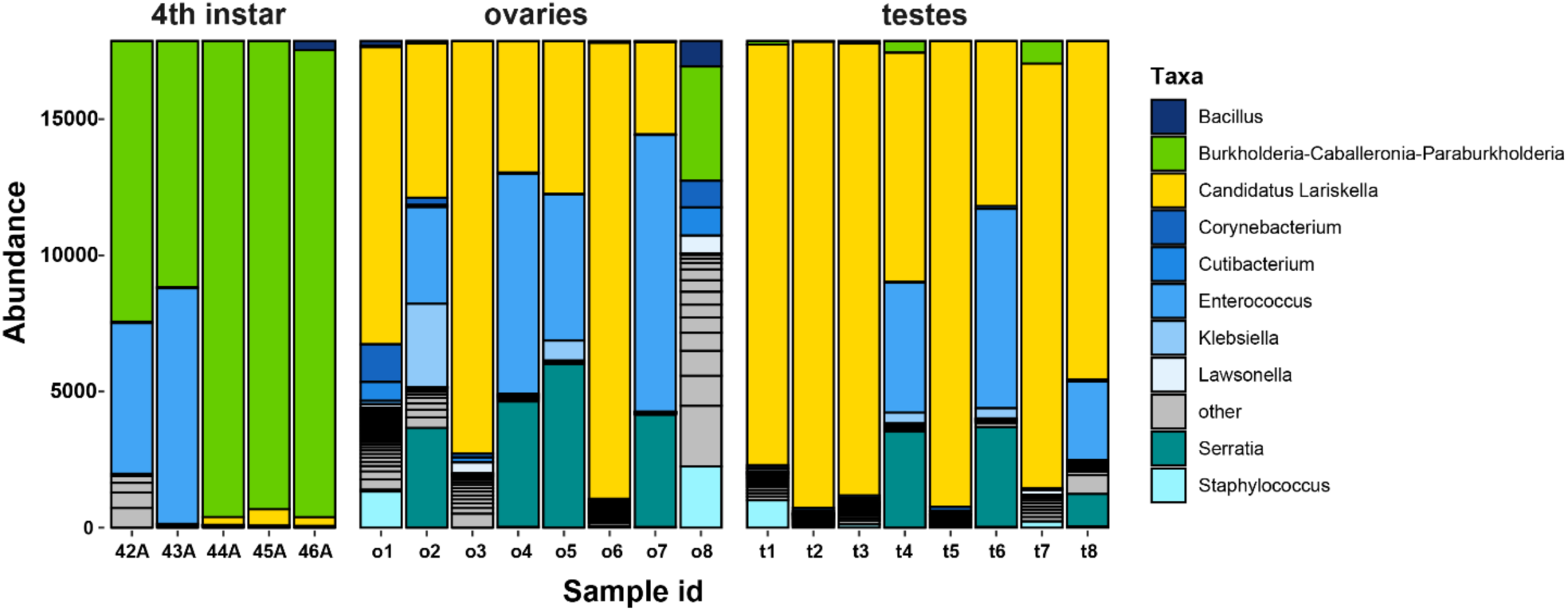
*Leptoglossus zonatus* amplicon 16S rRNA sequences from whole-body 4^th^ instar nymphs and reproductive tissue (ovaries and testes). Each bar represents an individual, and the colors represent reads of bacterial taxa denoted in the caption. In whole-body samples, the gut-associated bacterium *Caballeronia* was the most abundant followed by *Enterococcus* and *Lariskella. Lariskella* was the most abundant bacterium in the reproductive tissue followed by *Enterococcus* and *Serratia*.

### *Caballeronia* effects on performance and fitness of *Leptoglossus zonatus*

Pairwise comparisons of insect performance and fitness were conducted among *Lariskella* negative (L-) and positive (L+) and *Caballeronia* negative (C-) and positive (C+) bugs. The individuals in both *Caballeronia* negative treatments showed the major fitness deficits we expected from a previous study (53). Survival analysis showed a significant decrease in lifetime survival for bugs that did not receive *Caballeronia* (L-C-/L-C+, *z* = 6.60, adjusted *p* < 0.001 and L+C-/L+C+, *z* = 4.07, adjusted *p* < 0.001; Fig.3).

**Fig. 3.**
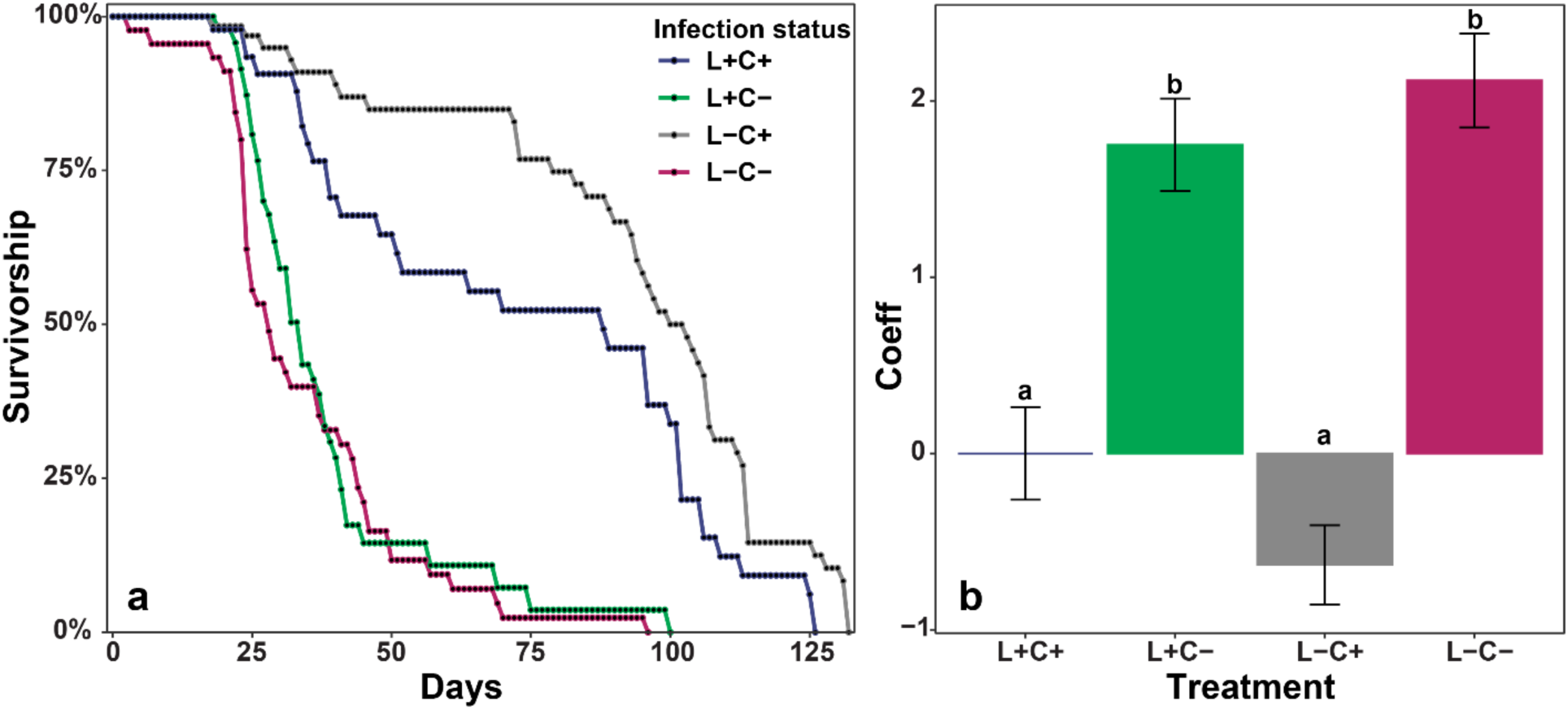
**a.** Kaplan-Meier survival curves showing the total lifespan of individuals from the 2^nd^ instar nymphal stage (when *Caballeronia* was acquired) based on the presence or absence of the primary symbiont *Caballeronia* (C+ or C-) and the presence or absence of the secondary symbiont *Lariskella* (L+ or L-). **Fig. 3b.** Coefficients of a mixed-effects Cox regression model in which higher coefficients indicate lower survivorship. The model shows a significant decrease in survivorship for bugs that lack *Caballeronia* regardless of *Lariskella* infection status. It also shows that *Lariskella* presence or absence does not significantly influence survivorship (L-C-/L+C-, adjusted *p* = 0.36 and L-C+/L+C+, adjusted *p* = 0.21, all other pairwise comparisons, adjusted *p* < 0.001).

The few survivors of the *Caballeronia*-negative treatments showed significantly longer development times (L-C-/L-C+, *t* = 6.55, df = 16.3, adjusted *p* < 0.0001; L+C-/L+C+, *t* = 7.42, df = 19.2, adjusted *p* < 0.001; Fig. 4). There was also no evidence that *Lariskella* was able to rescue bugs that lacked *Caballeronia*; the lengthened development times were equivalent in *Caballeronia* negative bugs with and without *Lariskella* (L-C-/L+C-, *t* = 1.557, df = 17.8, adjusted *p* = 0.42; Fig. 4). Although a few *Caballeronia* negative bugs did eclose as adults, females weighed significantly less than *Caballeronia* positive bugs (L+C+/L+C, *t* = 3.33, df = 27.3, adjusted *p* = 0.01 and L-C+/L- C-, *t* = 3.20, df = 27.7, adjusted *p* = 0.017; Fig. 5a). Similarly, *Caballeronia* negative males weighed significantly less than their *Caballeronia* positive counterparts, (L+C+/L+C-, *t* = 5.573, df = 27.58, adjusted *p* < 0.0001 and L-C+/L-C-, *t* = 3.72, df = 24.15, adjusted *p* = 0.0054; Fig. 5b). Lastly, when *Caballeronia* negative females were paired with mates, no female produced any eggs, indicating that *Caballeronia* is required for *L. zonatus* reproduction.

**Fig. 4.**
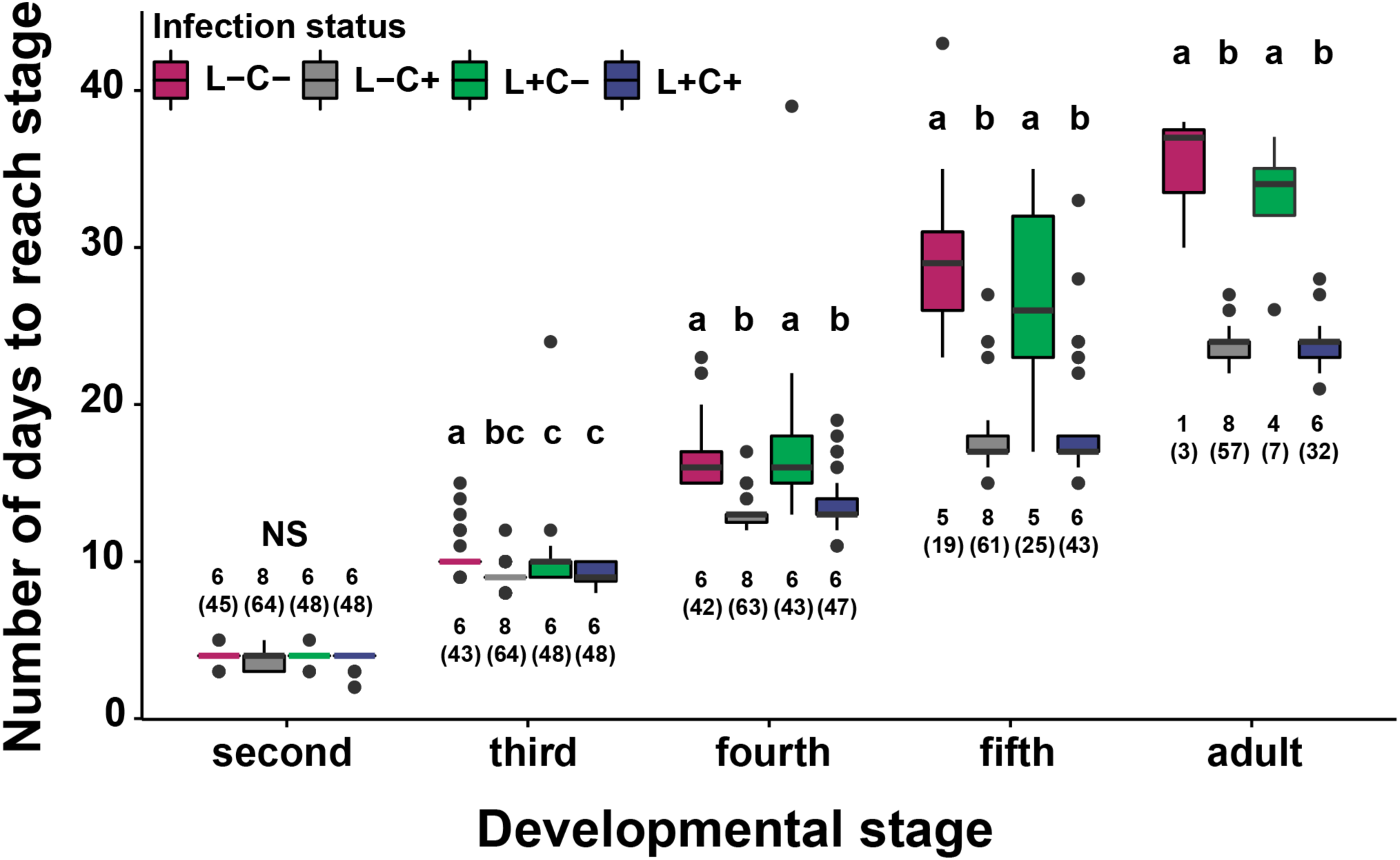
Development times for four treatments across developmental stages starting at the molt into the 2^nd^ instar when the bugs were fed either *Caballeronia* (C+) or deionized water (C-). Development time lagged significantly for the *Caballeronia* negative treatments but was not significantly different between *Lariskella* positive and negative bugs (adjusted *p* > 0.4 for *Lariskella* comparisons while keeping *Caballeronia* status constant). Bars with different letters reflect statistically significant differences. Numbers next to the bars indicate the number of replicates analyzed, with total numbers of individuals measured in parentheses.

**Fig. 5.**
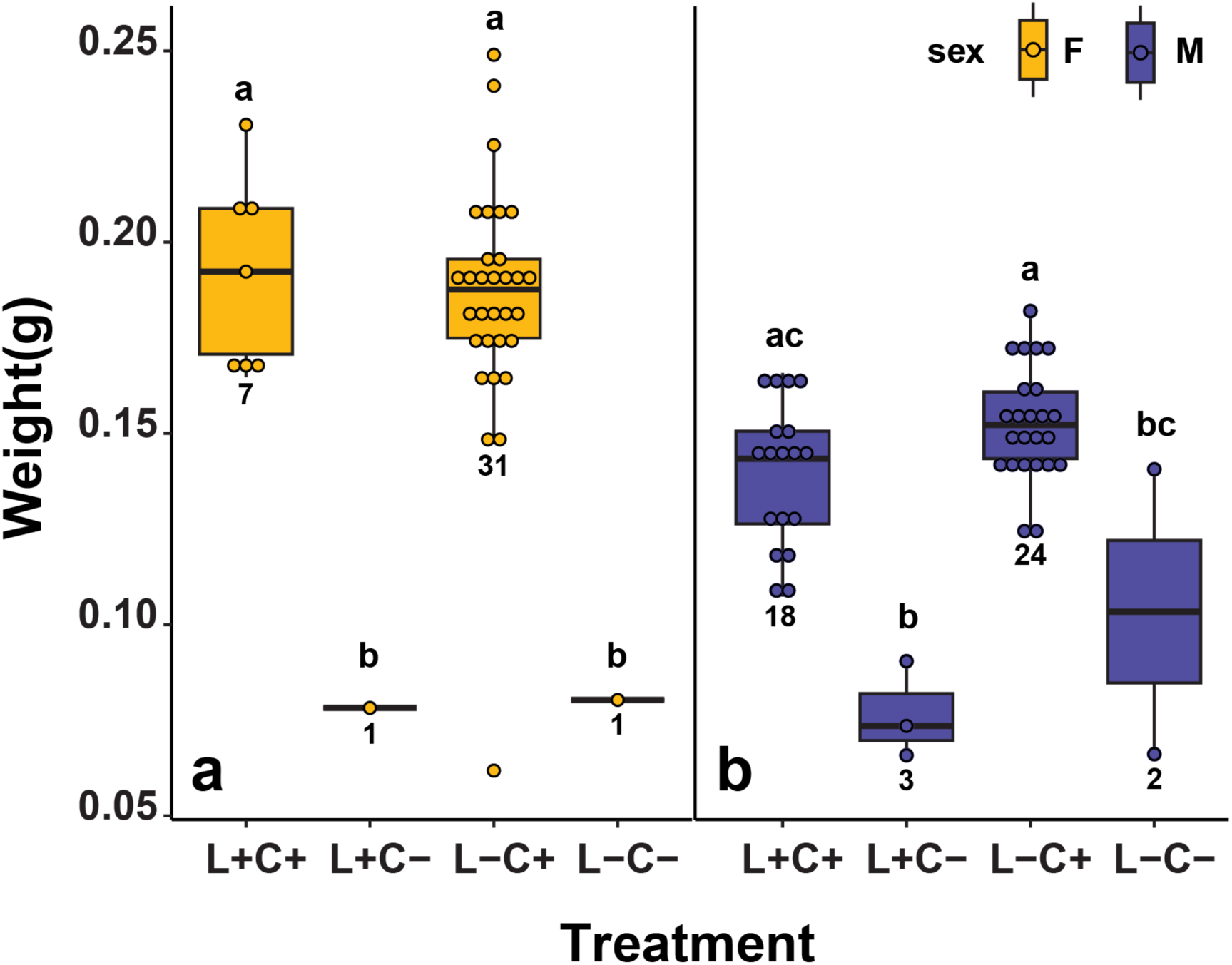
**a.** Mean weights of adults from the fitness experiment, showing a significant difference in weight between *Caballeronia* positive and negative females (C+, C-) regardless of *Lariskella* status (adjusted *p* > 0.9 for *Lariskella* comparisons while keeping *Caballeronia* status constant and all other pairwise comparisons, adjusted *p* < 0.018). **Fig. 5b.** Although there was a significant difference in weights among C+/C- males when controlling for *Lariskella* status (adjusted *p* < 0.01), there was no significant difference between L-C-/L+C+ males (adjusted *p* = 0.0571).

### *Lariskella* effects on performance and fitness of *Leptoglossus zonatus*

In contrast to the findings for *Caballeronia*, there were no significant differences in lifetime survival between the treatments with and without *Lariskella* (adjusted *p*-value of all possible *Lariskella* comparisons > 0.21; Fig. 3). Similarly, adults that were *Lariskella* positive developed at equivalent rates to individuals that lacked *Lariskella* (L-C+/L+C+, *t* = -0.226, df = 13.9, adjusted *p* < 0.99; Fig. 4). Finally, adult female weight was not influenced by the presence of *Lariskella* (L+C+/L-C+, *t* = 0.55, df = 14.0, adjusted *p* = 0.945 and L+C-/L-C-, *t* = -0.047, df = 29.4, adjusted *p* = 1.0; Fig. 5a). *Caballeronia* positive males with and without *Lariskella* were also similar weights (L+C+/L-C+, *t* = - 2.288, df = 9.93, adjusted *p* = 0.17; Fig. 5b).

Of the egg laying adults in the treatments with *Caballeronia*, paired adult females had a long reproductive period of about 100 days. Both clutch size and egg hatch rates declined throughout the life of the female, so time was a significant factor for both (Fclutch size = 25.08, df = 258, *p* < 0.001; Fig. 6a), (F hatch rate = 7.757, df = 258, *p* < 0.001; Fig. 6b). However, the fecundity of reproducing females was not influenced by the presence of *Lariskella* (*t* = 0.757, *p* = 0.45) with females producing approximately 300 eggs over their lifetime whether *Lariskella* was present or absent (Fig. 7). Similarly, the presence of *Lariskella* did not influence egg hatch rate (*t* = -0.389, *p* = 0.69; Fig. 6b).

**Fig. 6.**
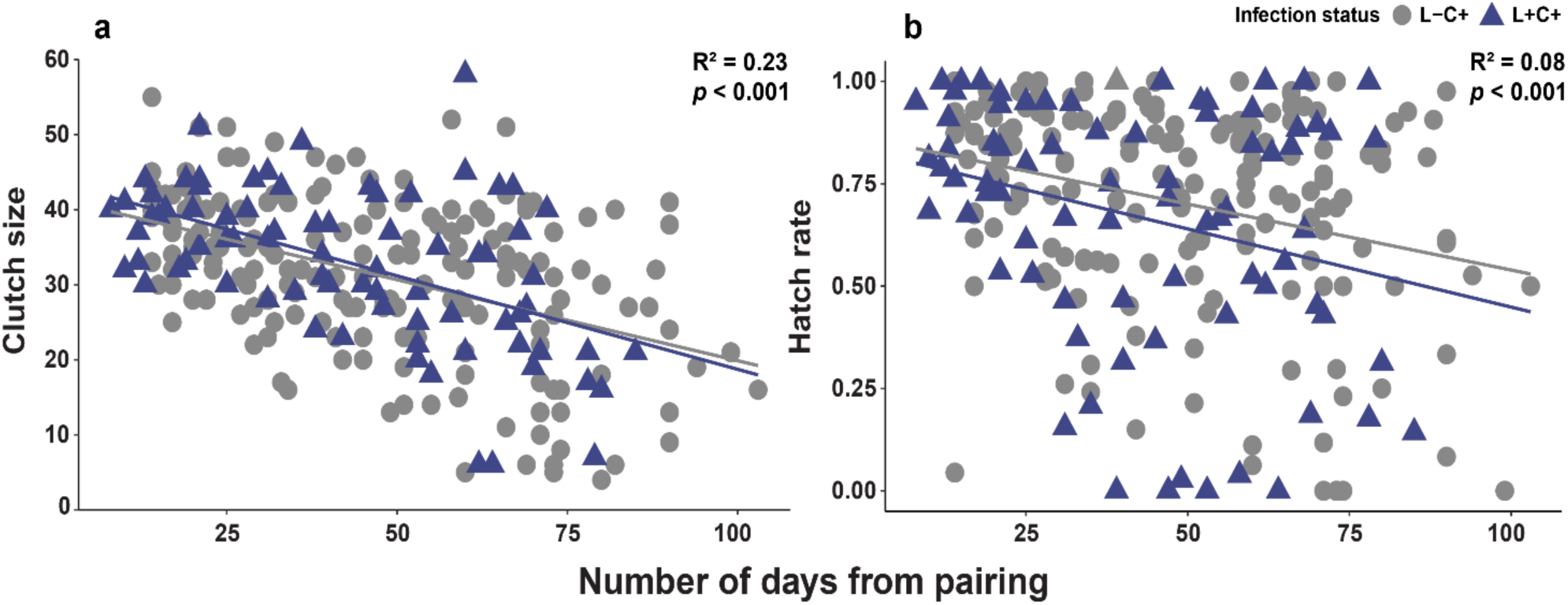
Scatterplots of **a)** clutch size and **b)** egg hatch rate of adult female *L. zonatus* with and without *Lariskella* over the lifetime reproductive period of ∼100 days. The fecundity of females (*Caballeronia-*positive treatments only) was not influenced by the presence of *Lariskella* (*p* = 0.45 and *p* = 0.70 for clutch size and hatch rate respectively). Both clutch size and egg viability (hatch rate) declined significantly throughout life.

**Fig. 7.**
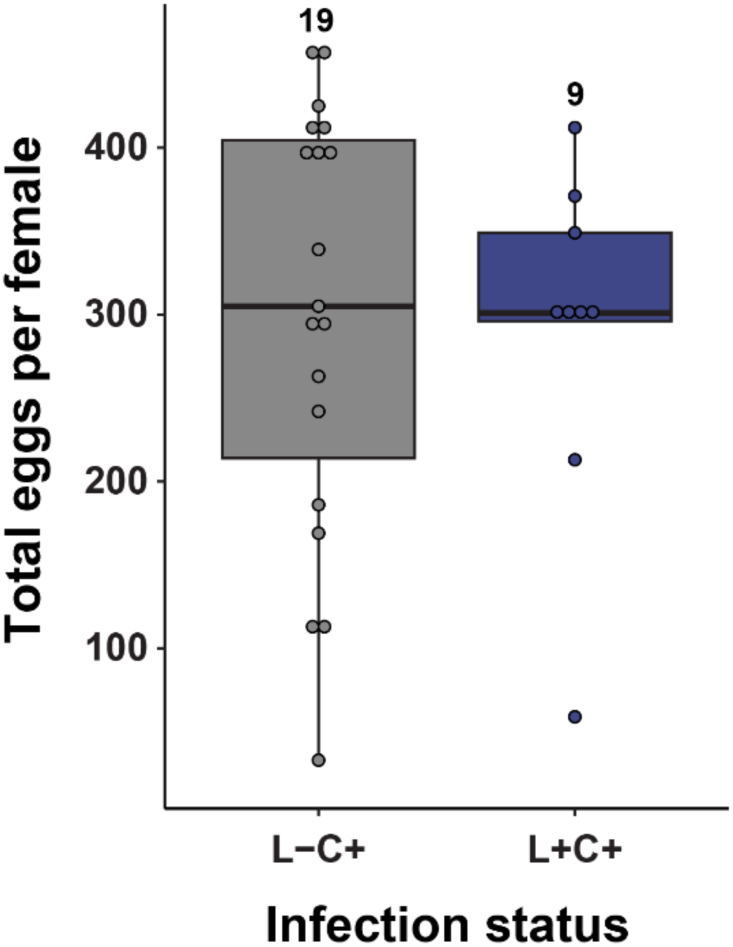
The presence of *Lariskella* did not influence the lifetime number of eggs produced by *Caballeronia* positive females during their lifetime (*t* = -0.23, *p* = 0.82). No *Caballeronia* negative adult females laid eggs, so those two treatments are absent from this figure.

### Cytoplasmic incompatibility (CI) crosses

In a second experiment, males and females with and without *Lariskella* were crossed in all four possible combinations to evaluate the possibility that *Lariskella* caused cytoplasmic incompatibility. In evaluating egg mortality in these crosses, we distinguished between early embryonic mortality which appeared to be relatively uncommon, and the frequent late embryonic mortality seen in offspring of aging females (e.g., Fig. 6b, and see Methods, Fig. 10). In the putative CI cross, with L+ males mated with L-females, there was a significant decrease in early embryonic survival of offspring relative to the other three crosses (χ^2^ = 23.44, df = 3, *p* > 0.0001, Fig 8), consistent with the pattern expected in a CI phenotype. Cytoplasmic incompatibility is not complete, but survival of eggs laid by females in the CI cross was less than two thirds that of eggs in the other three crosses.

**Fig. 8.**
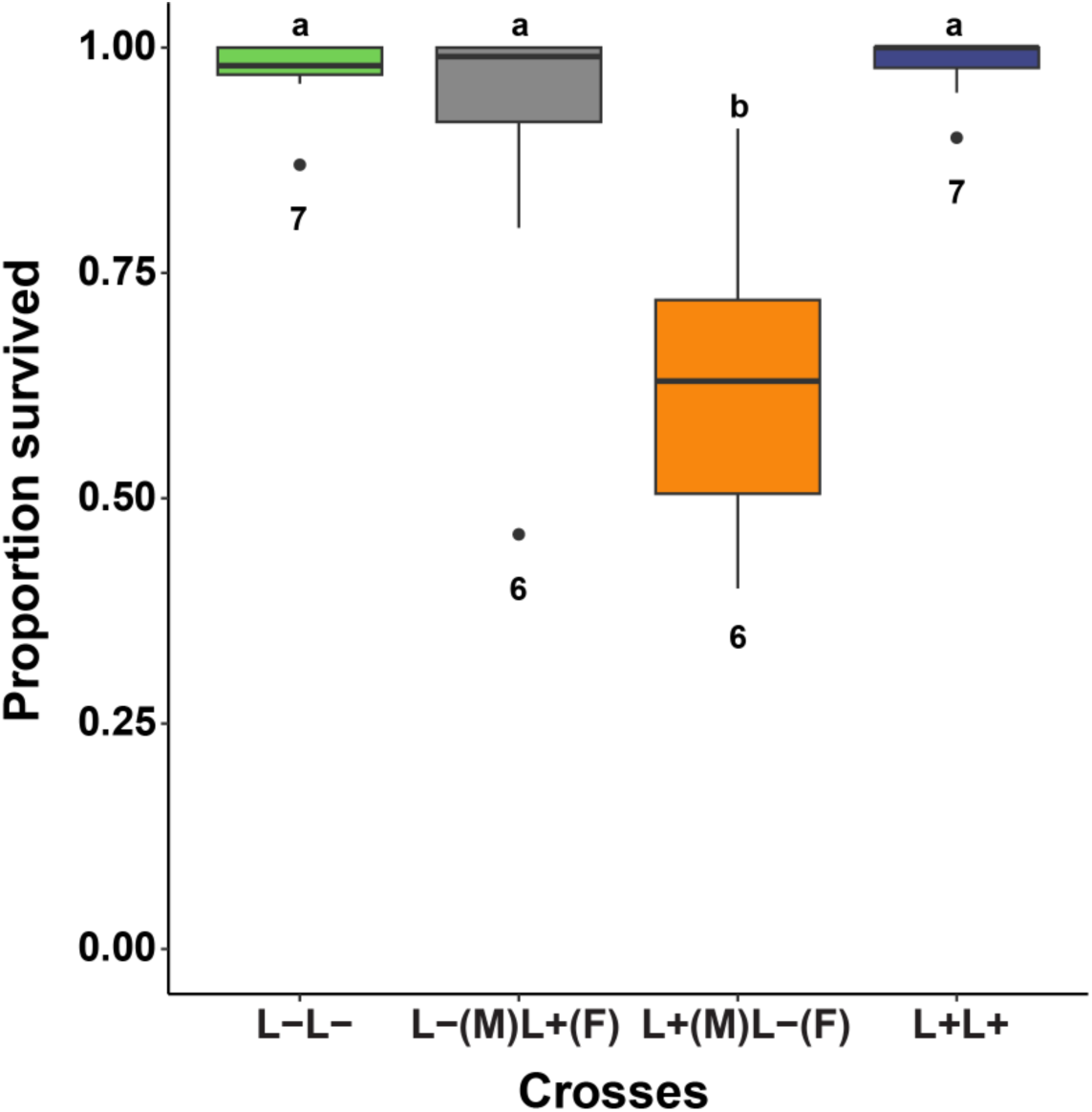
The proportion of a female’s eggs that survived early embryonic development among crosses of *Lariskella-*infected (L+) and uninfected (L-) adults in *Caballeronia* + *L. zonatus*. Significantly fewer eggs survived early embryogenesis in the putative CI cross than in any of the other crosses (*p* < 0.001), suggesting that *Lariskella* causes CI in *L. zonatus*. Numbers under bars refer to the number of replicates.

### *Lariskella* frequency in the field

The proportion of bugs infected with *Lariskella* was high but not fixed for *L. zonatus* samples collected in California and Arizona, USA (Fig. 9a). In a Tucson pomegranate orchard sampled repeatedly over two seasons, the proportion of *Lariskella* positive bugs in samples was variable and ranged from a low of 66% in July and August of 2019 to a high of 100% in June and July of 2020, but showed no clear seasonal pattern (Fig. 9b).

**Fig. 9.**
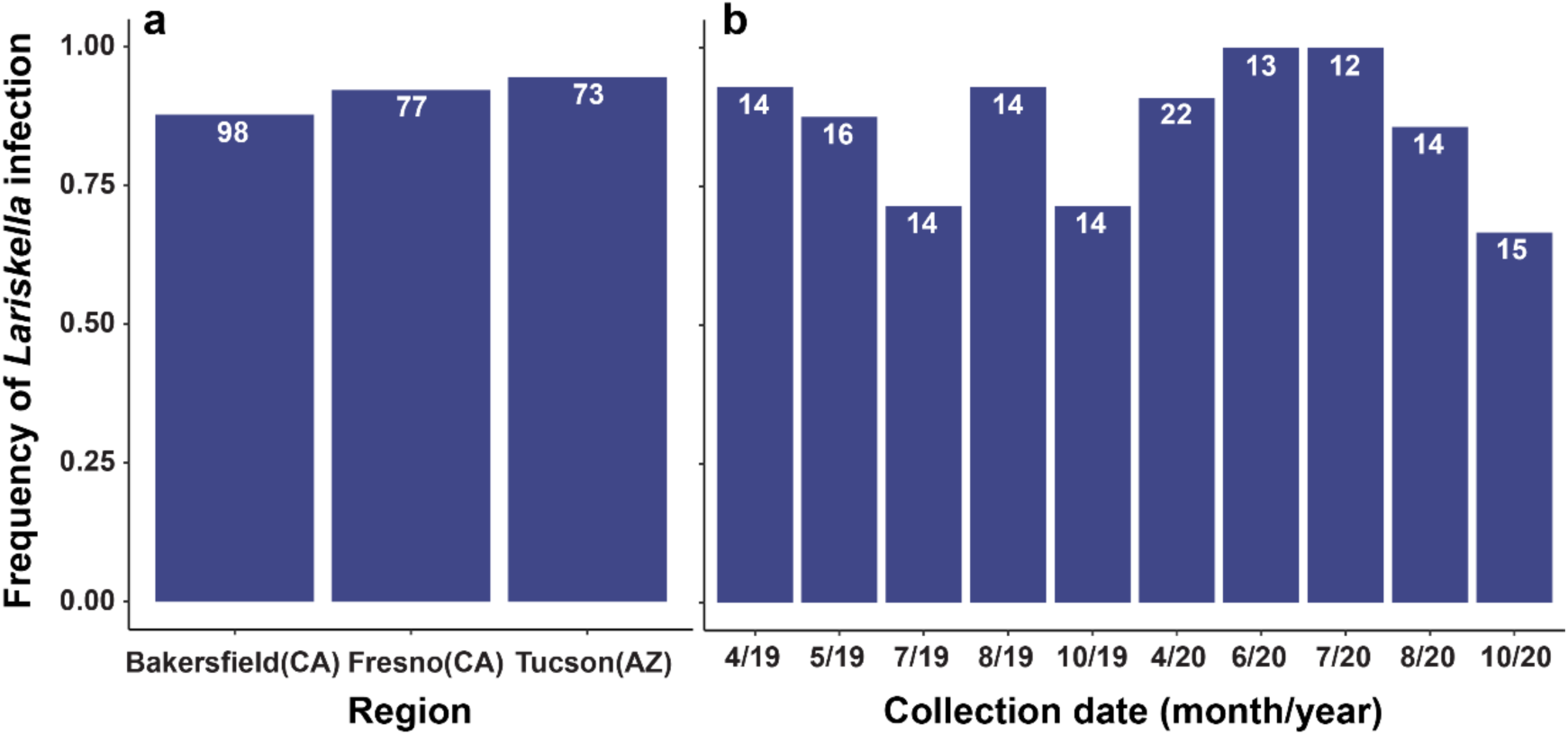
**a.** *Lariskella* infection frequencies in three *L. zonatus* populations in USA on pomegranates: two populations in California (Bakersfield and Fresno) and one from Tucson, Arizona. *Lariskella* frequencies were high (0.88 Bakersfield, 0.92 Fresno, and 0.94 Tucson, but were not fixed in any population. **Fig. 9b.** *Lariskella* infection frequency in *L. zonatus* over time in a single pomegranate orchard in Tucson, AZ, USA. Bugs were sampled at approximately 6-week intervals over the bugs’ active period from April to October in 2019 and 2020.

## Discussion

We examined the role of the intracellular symbiont *Lariskella* on *Leptoglossus zonatus* fitness when the obligate nutritional symbiont, *Caballeronia* was present or absent. We found no fitness costs or benefits to *Lariskella* throughout the lifetime of *L. zonatus*, nor did we find an interaction with *Caballeronia*. Crosses instead revealed that *Lariskella* causes incomplete cytoplasmic incompatibility (CI). High but not fixed frequencies of *Lariskella* were found in field populations. The high frequencies would be predicted for a symbiont that spreads via CI, has near-perfect maternal transmission and an absence of fitness costs (68–71).

In the current study, bugs with and without *Lariskella* had equivalent development times and lifetime survivorship. However, most individuals lacking the primary symbiont, *Caballeronia,* died before reaching adulthood. Although a few *Caballeronia*-negative individuals survived for several months, nearly all failed to mature, and all that did were unable to reproduce. This result provides even more support for an earlier conclusion that *Caballeronia* is obligate for *L. zonatus* (53). Our results further suggest that *Lariskella* cannot offset the severe fitness costs experienced by bugs lacking *Caballeronia*, at least under the laboratory conditions tested. This does not rule out a possible nutritional role for *Lariskella,* however. If signaling by the presence of *Caballeronia* in the gut is necessary for basic functions like gut development as has been found in *R. pedestris* (72), it could be that even if *Lariskella* synthesized all the nutrients limiting for *L. zonatus,* it would not compensate for the absence of the development regulating function of *Caballeronia.* Further, *Caballeronia* alone may provide all limiting nutrients in abundance such that any nutrient biosynthesis of *Lariskella* is entirely redundant. Nevertheless, it remains possible (though, we speculate unlikely) that *Lariskella* could confer a nutritional benefit that becomes evident when *L. zonatus* is paired with a suboptimal *Caballeronia* strain or close relative.

Previous work showed that some *Caballeronia* strains are more beneficial to *L. zonatus* development and adult weight than others (53), and allied genera in the Burkholderiacae can colonize the *R. pedestris* gut and provide some benefits but are inferior to *Caballeronia* for bug fitness (61). Perhaps a better test of a nutritional role for *Lariskella* would be to introduce a suboptimal *Caballeronia* that triggers the normal developmental program of the bug but falls short of providing complete nutrition for *L. zonatus*. In this situation, a nutritional role of *Lariskella* could benefit *L. zonatus*. Analysis of the genome of the CI-causing *Lariskella* in *L. zonatus,* when available, will allow us to predict whether *Lariskella* could complement the nutrition provided by a suboptimal *Caballeronia* or other Burkholderiaceae strain.

The recent emergence of *Lariskella* as a relatively common symbiont of arthropods underscores the mystery of the family to which it belongs, the Midichloriaceae. This family is the most diverse yet least understood within the intracellular bacterial order Rickettsiales, an order of alphaproteobacteria that includes significant human and livestock pathogens, with evidence suggesting a Rickettsiales was the ancestor of mitochondria (40,73–76). Members of Midichloriaceae are intracellular symbionts found across a wide range of hosts and habitats, predominantly aquatic.

They have been detected in unicellular protists (e.g., amoebas, ciliates) and invertebrates (e.g., ticks, corals, and arthropods), reflecting their complex ecological distribution among aquatic and terrestrial hosts. Despite a shared intracellular lifestyle, their genomes vary significantly in size and gene content, including differences in metabolic pathways even among closely related genera, which suggests frequent horizontal gene transfer and evidence of recent host shifts (40,77). The type genus *Midichloria* has so far been found exclusively in ticks, particularly *Ixodes* ticks, where some species inhabit the mitochondria (78–80). *Midichloria* appears to function as a nutritional symbiont, synthesizing folate, biotin, and B vitamins necessary to supplement the vertebrate blood diet of its tick hosts (78,81). Given the rampant horizontal gene transfer and hosts shifts in Midichloriaceae, the potential exists for certain strains of *Lariskella* to supplement nutrition in blood-feeding and non-blood-feeding arthropods, particularly for B vitamins which are often equally deficient in plant sap and blood (82,83). Demonstrating the functional roles of Midichloriaceae members in their hosts has proven challenging, partly because they are uncultivable and partly because of the potential difficulty of curing hosts using antibiotics, especially in cases where hosts rely on multiple obligate symbionts (84). The current study adds cytoplasmic incompatibility (CI) as another phenotype associated with Midichloriaceae.

*Wolbachia*, in the Anaplasmataceae family of Rickettsiales, was thought to be unique in causing CI for several decades after the phenomenon was first documented (13,14,20). Since then, representatives of five other bacterial lineages have been shown to cause CI, including two other Alphaproteobacteria (*Mesenet* & *Rickettsia)*, *Ricketsiella* (Gammaproteobacteria), *Cardinium* (Bacteroidota), and *Spiroplasma* (Mollicutes) (11,85–88). The current work may be the first characterization of the functional role of *Lariskella* in any host and places this bacterium among a group of now seven lineages that cause CI.

We do not know whether *Lariskella* causes CI in other hosts, but here the frequency of hosts carrying *Lariskella* may give some hints. Theory predicts that the invasion of a CI symbiont with a near perfect maternal transmission rate coupled with a lack of fitness costs should result in high frequencies or fixation in a population (68–71,89). Although *Lariskella* infection in *L. zonatus* was not fixed in any of the California or Tucson populations surveyed, all three sites had a similarly high frequency of *Lariskella* infection (>85%), similar to the numbers previously observed for *Nysius* seed bug species in Japan (46) and *Ixodes* ticks in Russia and Japan (41–43). These high frequency infections would be consistent with either CI or a nutritional role. Within *L. zonatus* populations, several factors could explain the lack of fixation in the field, including an incomplete CI phenotype and low symbiont titer early in host development that may render the symbiont vulnerable to environmental stressors like heat or environmental antibiotic exposure (90–92). Additionally, the long reproductive period of *L. zonatus* may reduce CI strength, as CI *Wolbachia* have been shown to decline with male age in some systems (93). While low prevalence of *Lariskella* in other arthropod species could suggest a number of scenarios including a relatively recent, horizontally acquired association, or one that is asymptomatic and slowly declining, it may also indicate a conditional role other than CI such as defense or temperature stress mediation (7,94).

Understanding *Lariskella*’s role in *L. zonatus* helps provide more insight into the biology and evolution of this insect. As one example, the mitochondrial population structure of *L. zonatus* in California was found to show evidence of positive selection or a bottleneck, with only three mitochondrial haplotypes compared to its sympatric congener *L. clypealis* with 17 (95). In light of the current study, low haplotype diversity in *L. zonatus* may be evidence of a relatively recent sweep of *Lariskella* that brought a coinherited mitochondrial haplotype along with it (96). The findings could also inform pest management strategies for tree crops and other hosts and may also provide insight into management of blood-feeding arthropods that vector human pathogens. Although the current study observed only a mild CI phenotype (with 40% of eggs affected), more examples of *Lariskella* CI strains and host backgrounds are needed to determine the range of CI strength that can be caused by this lineage. In *Wolbachia* and *Mesenet*, CI can cause complete (100%) offspring mortality (85,97–99), but CI strength in *Wolbachia* varies tremendously depending on both the symbiont and the host genotype (98,100). Recent work shows that incomplete? CI *Wolbachia* strength in *Culex pipiens* can be ameliorated by the divergence of CI gene repertoires relative to strains that induce complete CI (100). Conversely, in *Drosophila*, *Wolbachia* can produce CI strength that varies from 30% to 100% mortality, depending on host species (98). The identification of a new CI lineage, *Lariskella,* that also infects human disease vectors (e.g., ticks and fleas) is notable given the pathogen-blocking effects of CI *Wolbachia* in mosquitoes (37,101,102). CI *Wolbachia* pathogen-blocking has spurred a global program deploying *Wolbachia* in mosquitoes to combat RNA viruses responsible for deadly diseases (103).

Future research should focus on comprehensive screening of *Lariskella* across a broad range of arthropods and comparative genome sequencing to characterize its metabolic pathways, potential nutritional roles, and capacity to manipulate host reproduction via homologs of known cytoplasmic incompatibility (CI) genes. For example, *Mesenet*, another alphaproteobacterium within Rickettsiales, carries homologs to *Wolbachia* CI genes (85). Both *Mesenet* and *Wolbachia* belong to the family Anaplasmataceae, which phylogenetic analyses often identify as a sister group to Midichloriaceae. Given the widespread occurrence of horizontal gene transfer and host shifts, *Lariskella* may have evolved CI independently, similar to *Cardinium*, which lacks *Wolbachia* CI genes (104,105). Alternatively, *Lariskella* could represent a novel mechanistic model of CI that diverges from the toxin-antidote, and two-by-one models observed in *Wolbachia*. Understanding the prevalence of *Lariskella*, its evolutionary trajectory, and its interactions within arthropod hosts will advance our knowledge of symbiont-driven reproductive manipulation and vector ecology. This research could also provide insights for developing new control strategies for pest and pathogen vectors.

## Materials and methods

### Leptoglossus zonatus culture

*Leptoglossus zonatus* adults were collected at the West Campus Agricultural Center pomegranate orchard maintained by the University of Arizona (Tucson, AZ, USA) in 2018 and established in the laboratory in large, screened plexiglass cages (30 X 30 X 30 cm) in a walk-in incubator set at 27°C, 16L:8D. The cages contained whole cowpea plants *(Vigna unguiculata*) potted in PRO-MIX MP potting mix in 15 cm pots with raw Spanish peanuts glued to index cards for food.

### Generating *Lariskella*-free cultures

First and 2^nd^ instar nymphs were fed 75µl of rifampicin-saturated EtOH in 1 ml H2O. They were fed the antibiotic for 3 days, then given deionized water with 0.05% ascorbic acid (DWA) for 3 days before the nymphs were fed with *Caballeronia* for 2 days. Once these individuals reached adulthood and reproduced, a portion of the newly hatched offspring was sacrificed and screened for *Lariskella* 16S rRNA via diagnostic PCR using the Duron et al. 2017 primer set (Forward:MIDF2:CCTTGGGCTYAACCYAAGAAT) and (Reverse:LARISR2:TTCCCAGCTTTACCTGATGGCAAC). Individuals in cohorts with siblings that were all or mostly negative for *Lariskella* were kept in separate containers. These F1 1^st^ and 2^nd^ instar nymphs were then treated with a higher dose (150µl) rifampicin-saturated EtOH added to 1 ml H20, then fed *Caballeronia* as before and reared to adulthood and allowed to mate and lay eggs. Again, a portion of neonates from several egg clutches were tested and only individuals with siblings that were all or almost all negative for *Lariskella* were kept. These F2 1^st^ and 2^nd^ instar individuals were treated for one more generation with the higher F1 dose of rifampicin. After these three generations of antibiotic treatment, nymphs were tested from different clutches. Nymphs from clutches in which 100% of the tested individuals were found to be *Lariskella* negative were combined to produce the final *Lariskella* negative (L-) culture. The L-culture was maintained without additional antibiotics for > 50 generations in the same rearing room as the *Lariskella* positive (L+) culture and is periodically checked with diagnostic PCR to confirm symbiont status.

### Maternal transmission rate of *Lariskella*

To test for the maternal transmission efficiency of *Lariskella*, 113 eggs from 11 different females were collected, frozen at -20°C and individually extracted using the Qiagen DNeasy Blood and Tissue kit, and infection status was confirmed with diagnostic PCR and gel electrophoresis.

### Absolute *Lariskella* quantification throughout development and in reproductive tissue

To estimate *Lariskella* titer throughout development and in reproductive tissue, we reared a clutch of eggs from a single female and collected 4-6 individuals 2-4 days after hatching, as well as in each subsequent developmental stage. Collected individuals were stored at -80℃. Additionally, we isolated 8 pairs of testes and 8 pairs of ovaries from adult bugs within 48 h after eclosion and stored them at -80℃. Each set of testes and ovaries were snap-frozen with liquid-nitrogen, pulverized with a disposable pestle and DNA extracted using the Qiagen DNeasy Blood and Tissue kit. For whole-body insect DNA extractions, we followed the same method, but for 4^th^ instar to adult stages, we split individuals among 2-5 spin columns and combined the extractions after elution. This ensured that a maximum of 50mg of tissue homogenate was used per column as per manufacturer’s instructions.

Total DNA was quantified using the Qubit dsDNA assay and all extractions were kept at -20℃. To quantify exact *Lariskella* genome copies in *L. zonatus* individuals, a 1.3 kb of the single copy *dnaA* gene of *Lariskella* was amplified by TaKaRa ExTaq DNA polymerase by using *Lariskella* specific-primers designed by the available genome sequences of *Lariskella*, namely LardnaA_23F: TAGTTGATGTTGAGTCTCAT and LardnaA_1355R ACACTAGAATTATCGCTAAT, and the products were sub-cloned into a pt7Blue T-vector (Novagen). DNA sequences of sub-cloned fragments were further determined by BigDye terminator v3.1 cycle sequencing kit and ABI 3130xl genetic analyzer. Then, quantitative PCR was performed using *L. zonatus Lariskella* specific primers targeting the *dnaA* gene (Forward: LzLar_1113F: ACCTTCTATTACTGCAATAC) and (Reverse: LzLar_1216R: GCCTAGCAAGCACAGACTTTCC). A 3-step qPCR was performed with an annealing temperature of 54℃ for 40 cycles with an additional melt curve step using the Bio-Rad CFX Connect system using the Maxima SYBR Green Master Mix. The absolute *Lariskella* abundance was estimated using 10-fold serial dilution standards (10^8^-10^3^) made directly from the pT7-blue vector containing the cloned *Lariskella dnaA* gene. All standards, unknown DNA samples and negative controls were run in triplicate.

### Reproductive tissue and whole-body 16S Illumina sequencing

To verify that other reproductive manipulators were not present in *L. zonatus* (e.g. *Wolbachia, Cardinium*), 16S Illumina amplicon sequencing was performed on 5 whole-body 4^th^ instar nymphs, and 8 pairs of testes and ovaries. (These were the same samples collected for qPCR, described above.) Briefly, we followed Illumina’s two-step amplification protocol (106). In an initial PCR, we amplified the V3-V4 hypervariable regions of the 16S rRNA gene using primers 341F/785R (107). In a second PCR we added 8 bp barcodes to the forward and reverse ends of the amplicons; these uniquely identified each sample, allowing multiplexing. We sequenced an equal mass of each sample’s PCR product on a 600 cycle paired-end Illumina MiSeq run at the University of Texas Arlington’s Life Science Core Facility. A DNA extraction blank was included with and processed identically to the samples, including sequencing.

Adapters and primers were trimmed with cutadapt (108). Poor quality reads were removed and bacterial amplicon sequence variants (ASVs, which approximate bacterial strains) were inferred using the R DADA2 package (109). We performed *de novo* chimera checking and removal. Taxonomy was assigned using the RDP classifier with the SILVA nr99 v138 database as the training set (110,111). We removed mitochondria, chloroplasts, and reads that did not fall within the expected length of the amplicon (398-445 bp). Contaminants were identified via the R decontam package’s *isContaminant* function, using a stringent threshold of 0.5 (112). This resulted in removal of four contaminants belonging to the genera *Acinetobacter, Ralstonia, Rahnella,* and *Micrococcus*. Data were rarefied to 17875 reads per sample, where sample rarefaction curves had plateaued.

### *Lariskella* fitness effects and interaction with *Caballeronia*

Insects were reared with and without *Lariskella* and with and without *Caballeronia* in a factorial design to test the effects of *Lariskella* on fitness, as well as the possibility of an interaction between the intracellular *Lariskella* and nutritional gut symbiont *Caballeronia*. We measured insect developmental mortality, development time, weight at adulthood, lifespan, lifetime fecundity, and egg viability (hatch rate). We reasoned that if *Lariskella* had a nutritional role we might expect greater fitness of *Caballeronia*-negative bugs when *Lariskella* was present.

### Insect rearing and *Caballeronia* feeding

Eggs were collected from both *Lariskella* positive (L+) and *Lariskella* negative (L-) *L. zonatus* cultures. The eggs were transferred to Petri dishes supplied with water tubes (containing DWA). After confirming *Lariskella* status, early 2^nd^ instar nymphs (the first feeding stage) were distributed into 16 L+ boxes and 16 L-plexiglass boxes (11.33 cm x 11.33 cm x 4 cm) with mesh lids. Eight nymphs were placed into each box and provided with raw peanuts for food, but initially, no water. Twenty-four hours later, the nymphs in eight of the L+ boxes and eight of the L-boxes were fed an aqueous suspension of *Caballeronia* cells (10,000 cfu/µl) once per day for 3 days. The remaining 8 L+ and 8 L-boxes were fed water alone and served as *Caballeronia-*negative treatments. After the third day of *Caballeronia* or water-only feeding, water vials were returned to all boxes and a single cowpea (*Vigna unguiculata*) seedling in a tube with water agar was added to each box. Seedlings were replaced as needed and water vials refilled until the bugs reached adulthood.

### Tracking insect development and adult fecundity

Nymphal development and mortality were tracked daily until adulthood. Bugs that died before reaching adulthood were removed from the rearing box and the date of death and development stage was recorded. The fresh weight of each adult was measured within 48 h after eclosion and then adults were paired within each of the four treatments (L-C-, L-C+, L+C-, L+C+). The pairs were placed in small cages (transparent, lidded plastic 500ml drink cups), each with peanuts, a water vial, and a single cowpea seedling. The pairs were monitored daily until the female died. When males died, they were replaced by other males from the same treatment (8 males replaced in total). Each day, egg clutches were collected from the cups and transferred to individual Petri dishes where eggs were counted and hatching success was measured.

### Survival, development, weight and fecundity analysis

To assess the effect that infection status (L-C-, L-C+, L+C-, L+C+) had on *L. zonatus* lifespan, survivorship was analyzed using a mixed-effects Cox regression model using the coxme R package (113) with modified R code from Duarte et al. (114) and cage as a random effect. The effect of treatment on time to reach each developmental stage was analyzed using a mixed-effects generalized linear model, with time (number of days to reach each stage post-*Caballeronia* feeding), and the presence or absence of *Lariskella* and *Caballeronia* as explanatory variables and cage as a random effect using the R package lme4 (115). The effect of infection status on adult weight was also analyzed using the same mixed-effects model for males and females separately with adult weight as the response variable. Post-hoc multiple comparisons were done with the emmeans package (116) for survivorship, development time and weight resulting in adjusted *p*-values for these analyses.

Additionally, we analyzed the effect of *Lariskella* on lifetime fecundity of females. The few *Caballeronia* negative females that survived to adulthood failed to produce any eggs and were therefore excluded from analysis. The effect of *Lariskella* on lifetime reproduction of females was analyzed for the response variables total egg number, clutch size, and hatch rate, using a multiple linear regression model in R using the base stats package (v4.2.3, R Core Team 2023) with time (days from pairing) and the presence or absence of *Lariskella* as explanatory variables. The effect of *Lariskella* on total egg production was determined using a one-way analysis of variance (ANOVA) (v4.2.3, R Core Team 2023).

Data involving bugs in six of the 32 boxes (4 C-, 2 C+) were excluded from analysis because diagnostic PCR indicated *Caballeronia* was either acquired from contamination sometime during the experiment (4 boxes in C-treatments) or was not acquired during exposure to *Caballeronia* (2 boxes in C+ treatments) using the same methods used in (53). Data from these boxes were excluded because lack of *Caballeronia* acquisition has severe negative fitness effects (53), and late *Caballeronia* acquisition in boxes that were not supposed to have it would have had unknown effects on fitness.

### Cytoplasmic incompatibility (CI) crosses

When no apparent effects of *Lariskella* on *L. zonatus* fitness were found, the possibility of *Lariskella* causing cytoplasmic incompatibility was evaluated. If *Lariskella* caused CI, we would expect few or no eggs to hatch in the cross in which L+ males were mated with L-females.

*Leptoglossus zonatus* bugs were reared and fed using the same protocol described above, but in this experiment, all adults were *Caballeronia* positive. They were paired in all four possible crosses among *Lariskella* infected and uninfected bugs (L+female/L+male (n=7), L-female/L-male (n=7), L+female/L-male (n=6), L-female/L+male (n=6)). Eggs were collected from each pair at daily intervals for two weeks and held in Petri dishes for two weeks to monitor hatching.

Careful observation of offspring eggs showed two types of hatching failure. Unhatched pale, homogeneously-colored eggs appeared to have died early in embryogenesis (“early mortality;” Fig. 10a-b), while dark eggs often showed a well-developed embryo through the semi-transparent chorion that failed to eclose or died during emergence (“late mortality;” Fig. 10a-c). 1). Eggs were categorized into successful hatch, early-death and late-death embryos (Fig. 10). Late mortality of eggs appears to be common; we noticed these dark eggs in every treatment in the fitness experiment.

**Fig. 10.**
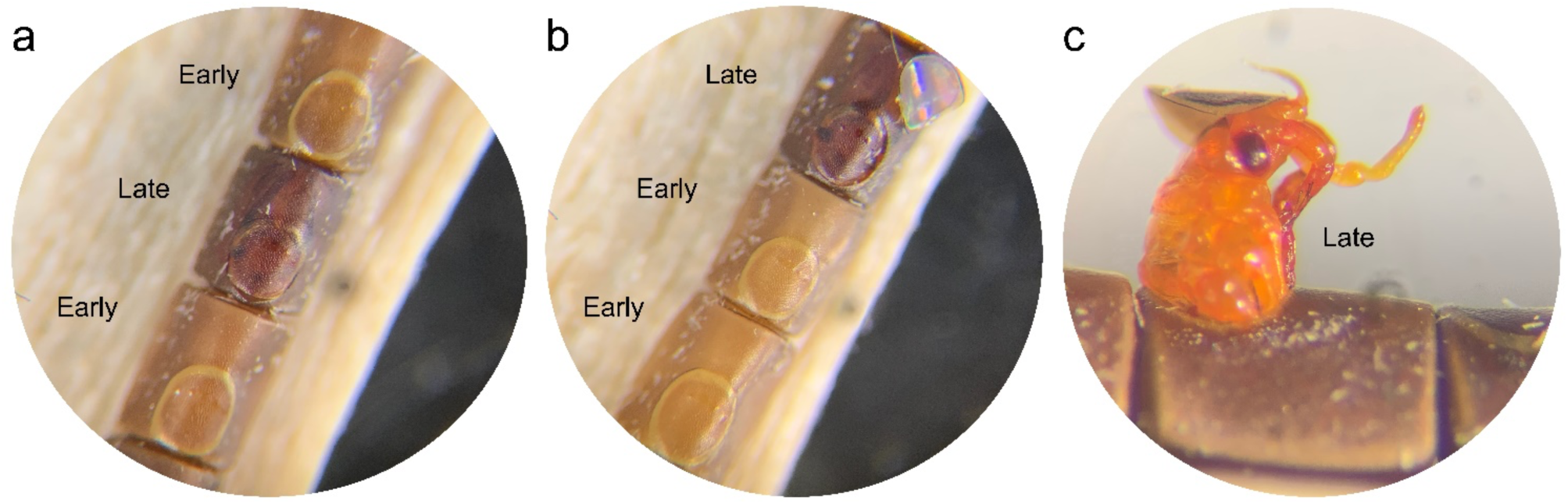
**a-b)** Unhatched *L. zonatus* eggs showing the pale homogenous color of eggs that died early in development (“early mortality”) and the dark brown color of eggs in which the embryo is well developed (“late mortality.”) **c**) Nymphs that died during emergence were included in the “late mortality” category.

We also hypothesized that CI would cause early embryonic mortality based on observations from other CI – inducing bacteria (15,117). To test for CI, we therefore compared exclusively early mortality among treatments, using the Kruskal-Wallis one-way analysis of variance and the Dunn pairwise test for pairwise comparisons between groups.

### Survey for *Lariskella* in field-collected *L. zonatus*

The lack of a performance or fecundity cost for *L. zonatus* bearing *Lariskella*, coupled with near perfect maternal transmission and moderate CI would all lead to a prediction that *Lariskella* in field populations of *L. zonatus* should be at high frequencies, but not likely fixed (118,119). To test this prediction, we surveyed *L. zonatus* adults collected from pomegranate in two locations in California, USA (Fresno and Bakersfield) and in Tucson, Arizona in 2018 (Ravenscraft et al, in review). All samples were stored in 95% ethanol. We also examined the frequency of *Lariskella* over time at one location, a University of Arizona pomegranate orchard adjacent to the Arizona Veterinary Diagnostics Laboratory, Tucson, AZ. A sample of adults was collected at approximately 6-week intervals from April – October in both 2019 and 2020.

### DNA extraction and diagnostic PCR to determine *Lariskella* infection in the field

Adult and nymphal bugs from the CA and 2018-2019 AZ samples were dissected, and DNA was extracted from a small portion of the M4 midgut region and surrounding tissue using the Qiagen DNeasy Blood and Tissue kit. DNA of bugs from the Tucson, AZ samples of 2020-2021 were extracted with an alternative, equivalent method. Abdomens were removed from the bugs, and entire abdomens were homogenized via bead beating. DNA extractions were performed with a small amount of the homogenate (5ul) and a Chelex extraction protocol (53). All the extractions were kept at -20℃ until diagnostic PCR was performed using *Lariskella* specific primers (42).

## Acknowledgments

We sincerely thank David Haviland for substantial advice and help in the field, the Kern County UC Cooperative Extension for use of their lab facilities, and Johnathan Adamson, Reiko Sekine, Chiaki Matsuura, and Alex Lombard for laboratory assistance.

We also thank White Forest Nursery (Bakersfield, CA), the Kearney Agricultural Research and Extension Center (Fresno, CA), the Mission Garden (Tucson, AZ), Mesquite Valley Growers (Tucson, CA), and Ursula Schuch (University of Arizona West Campus Agricultural Center pomegranate orchard) for permission to sample insects.

## Data Availability Statement

The datasets generated for this study can be found in the Dryad repository https://doi.org/10.5061/dryad.bvq83bkkp.

